# The small GTPase Rab11F represents a molecular marker within the secretory pathway required for the nitrogen-fixing symbiosis

**DOI:** 10.1101/661835

**Authors:** Prachumporn Nounurai, Holger Densow, Hanna Bednarz, Karsten Niehaus

## Abstract

The nitrogen-fixing root nodule is generally derived through a successful symbiotic interaction between legume plants and bacteria of the genus *Rhizobium*. A root nodule shelter hundreds of *Rhizobia*, which are thought to invade into the plant cells through an endocytosis-like process despite the existence of turgor pressure. Each invading *Rhizobium* is surrounded by the peribacteroid membrane to form the symbiosome, which results in the higher acquisition of host membrane materials. In this study, we show the localization of Rab11F, a RabA6b homolog with the large Rab-GTPase family, which was highly expressed in root nodules of *Medicago sativa* and *M. truncatula*. Rab11F-labeled organelles accumulated the membrane specific dye FM4-64 and were sensitive to Brefeldin A by forming aggregates after treatment with this drug. By co-localization with the cis-Golgi marker, GmMan1-mCherry, Rab11F-organelles formed tri-colored organelles, whereby Rab11F was located to the opposite side of GmMan1-mCherry indicating that Rab11F-labeled structures were localized within the trans-Golgi network (TGN). In root nodules, Rab11F was localized transiently at the infection thread-covering membrane on the side of infection droplets and the peribacteroid membranes. The symbiosome acquires Rab11F during the entry process and differentiation. However, the symbiosome did not recruit Rab11F after cessation of division. In conclusion, the *legume* plant seemed to use a specialized secretion pathway from the TGN, which was marked by Rab11F, to proliferate the symbiosome membrane.

## Introduction

Legume plants are able to live in symbiosis with *Rhizobium* bacteria resulting in the formation of specialized nitrogen-fixing root organs, termed root nodules. The onset of nodule development is triggered by the molecular signal exchange between the plant and the bacteria in which the bacterium produces lipochito-oligosaccharides, Nod factors, in response to the flavonoids secreted by the plant(1). In Turn, the Nod factor induces cellular responses in root hair cells and the reactivation of the cell cycle in the root cortex and the pericycle, leading to root hair curling and the formation of polarized cytoplasmic bridges (pre-infection threads) and a nodule primordium in the inner cortex (2,3).

Following a series of recognition stages, the bacteria become entrapped in the center of a root hair curl known as the hyaline spot, from which the bacteria enter the root via a tube-like structure termed infection thread. The infection thread is initiated by the invagination of the plant cell wall in the curl and by the degradation of the cell wall followed by the cell membrane invagination and the deposition of cell wall material around the distal end of the invagination. The lumen of the infection thread is topologically equivalent to the apoplastic, or the intercellular, space and is bordered by the infection thread wall and the membrane(2). It is filled with bacteria and a specific extracellular matrix of plant and microbial origin(4). When the infection thread reaches the nodule primordium, the bacteria are released into the host cell cytoplasm producing infection droplets, formed from the non-walled outgrowths of the infection thread. At the periphery of the droplet, the infection thread membrane adjacent to the bacterium invaginates and is subsequently pinched off, resulting in the deposition in the cytoplasm of a single bacterium surrounded by a lipid bilayer membrane, which is now termed bacteroid or symbiosome membrane(2). This process seems to be comparable to phagocytosis in animal cells.

The detailed mechanism of the endocytic process in plants has not been completely elucidated. In general, endosome-engulfed cargos are delivered to lytic vesicles via a series of endocytic compartments, beginning with the early endosome and then the late endosome, before finally being deposited in the lysosome in animals or the vacuole in plants(5,6). These changes in organelle characteristics can be identified by the presence of organelle-specific markers, viz. membrane receptors, enzymes, and small G-proteins, including Rab proteins(7). Rabs are small GTPases, which function as regulators of a variety of intracellular vesicle trafficking processes, such as endocytosis, exocytosis, and membrane recycling(8). Each Rab protein is located on a different intracellular membrane compartment. Thus, they can be used as organelle-specific markers. In animal cells, the compartment containing freshly engulfed bacteria is referred to as the early phagosome and contains the marker Rab5; subsequently, the phagosome recruits Rab7, a marker of the late endosome(9), which occurs by the removal of Rab5 with concomitant replacement by Rab7 (Rab conversion) (10,11).

In plants, symbiosomes do not acquire Rab5 at the early phagosome stage, although they recruit Rab7 when they stop dividing(12), suggesting that the symbiosome enters the host cytosol through a Rab5-independent endocytic pathway. However, localization of plant Rab5 homologs, namely Ara6/RabF1, Ara7/RabF2b, and Rha1/RabF2a is on multivesicular bodies (MVBs) or pre-vacuolar compartments, which are considered to be late endosomes(12–15). This suggests that symbiosomes are possibly formed by another uncharacterized endosomal compartment in plants.

There have been reports indicating that in plants, the trans-Golgi network (TGN) is involved in the formation of early endosomes(15–17). In *Arabidopsis*, VHA-a1-antibodies label organelles identified as TGNs, which rapidly internalize the endocytic tracer FM4-64 indicating that TGN and the early endosome (EE) are subdomains of the same compartment(16). In tobacco BY-2 cells, SCAMP1-labeled organelles are also TGN and absorb FM4-64 before the formation of the multi-vesicular body (MVB)/ the prevacuolar compartment (PVC)(15). In *Arabidopsis*, Rab-A2/A3 is localized to the organelles sensitive to the fungal toxin Brefeldin A, which accumulate FM4-64 in the vicinity of the prevacuolar compartment(17). The TGN is also involved in the formation of the cell plate during cell division, which is of special interest since the endocytosis of apoplastic material is also involved in the cell plate formation(18). In animals and yeast, the TGN acts as a recycling compartment in which Rab11 proteins are involved in membrane trafficking between the endosome, the TGN, and the plasma membrane(19,20). In polarized cells, the TGN also acts as the main sorting hub for directing secretory vesicles to the appropriate surface membrane destination (apical or basolateral)(21), a process regulated by Rab11(22). Therefore, the TGN has been proposed to be a specialized organelle with different subdomains responsible for directing vesicles to the lysosome for catabolism or to the plasma membrane for secretion(15).

Several studies have shown that plant Rab11 is associated with the secretory pathway from the TGN to the plasma membrane. In *Arabidopsis*, Rab11 or the RabA clade is the largest family among Rab families and has 26 members. There is certain evidence indicating that some members of the Rab11 family are involved in the symbiosis. In the symbiosis-defective mutant *dnf1* of *M. truncatula*, defective in a subunit of signal peptidase complex, the bacteroid and symbiosome development are blocked. The microarray analysis of *M. truncatula* gene expression atlas showed the expression of Rab11B highly correlated to the gene expression of DNF1(23). In the common bean (*Phaseolus vulgaris*), RabA2 RNA interference (RNAi) expressing plants failed to induce root hair deformation and the initiation of infection threads(24). Whitehead and Day (1997) already provided evidence for the origin of the symbiosome membrane. However, the molecular mechanism that determines membrane identity in this process remains unknown(25). Since small GTP-binding proteins are associated with specific membranes, these proteins can be used to distinguish between the otherwise microscopically identical vesicles. For this reason, these proteins are ideal tools for analyzing the origin of the symbiosome membrane.

In this study, the role of MsRab11F (Rab11F) in the symbiosome formation in *M. truncatula* nodules was investigated. Rab11F is highly expressed in *Medicago sativa* root nodules, which is consistent with the high proliferation of the symbiosome membrane, suggesting the involvement of Rab11F in this process(26). We show that Rab11F is transiently localized to the infection thread and the peribacteroid membranes, indicating that symbiosomes can recruit Rab11F immediately following the endocytic uptake process and during their development within the host plant.

## Materials and methods

### Construction of MsRab11f1-mGFP6 fusion protein

Msrab11f1 (Rab11F; accession number: AJ697970) was amplified by PCR from pFlag-Mac-Rab11F(26) using primers extended with BamHI and SacI restriction sites (<u>GGATCC</u>ATGGA-TCATGATGCAATTA, and <u>GAGCTC</u>TCATGAACAACAAGG AGCC) and was subcloned into pGemT-easy (Promega). The subcloned amplicon was digested with BamHI and SacI and ligated to the BamHI-SacI digested expression vector pET24a(+)(Novagen). The mgfp6 was amplified by PCR from p35S-<u>m</u>GFP6 using primers extended with BamHI restriction sites (<u>GGATCC</u>ATGCATAAAGGAGAAGAACTTTTCACTGG, and <u>GGATCC</u>TCACC CATCCTTTTTGTATAGTTCATCCAT) and subcloned into BamHI-digested pGemT-easy. As the C-terminal of Rab11F is needed for isoprenylation, which is essential for the attachment to the membranes(26), mGFP6 thus was fused to the N-terminus of Rab11F. The fusion gene was expressed under the control of the 35S promoter. The subcloned amplicon was digested with BamHI and inserted into the BamHI digested pET24-MsRab11f1 resulting in the pET24-mGFP6:MsRab11f1 expression vector, which was transfected into the *E. coli* strain BL21(DE3) and transformants were selected on LB medium plates containing the appropriate drug selection marker. In addition, positive colonies were identified by the presence of green fluorescence under UV-light.

To generate the vector for the transient transformation, we isolated the insert from the pET24-mGFP6:MsRab11f1 vector by digestion with XbaI and SacI and ligated into XbaI-SacI digested with p35S-EGFP (Clontech), thereby removing the wild type GFP.

To generate a construct for stable transformation, we amplified a sequence of MsRab11F-mGFP6 with the 35S promoter and NOS terminator coding sequences by PCR and subcloned into pGemT-easy. The amplicon was digested with HindIII and ligated into the HindIII digested binary vector pBIN19. The Msrab11f1(S29N) mutant was generated using QuikChange II Site-Directed Mutagenesis Kits performed according to the manufacturer’s manual (Stratagene) with the primers CTGGAGTTGGGAAAAACAATCTGCTTTCAAGG, and CCTTGAAAGCAGATTGTTTTTCCCAACTCCAG. The resulting binary vector was introduced into the *Agrobacterium rhizogenes* strain ARqua1 and the *Agrobacterium tumefaciens* strain GV3101:pMP90 by electroporation followed by the selection on YEP medium containing the appropriate antibiotics.

A 35S–soybean mannosidase I–red fluorescent protein (Gm-ManI-mCherry) construct was kindly provided by Dr. A. Staehelin and A. Nebenfuhr (University of Colorado, Boulder CO, USA)(27).

### *Medicago truncatula* growth, transformation, and nodulation

*M. truncatula* cv. Jemalong seed surfaces were sterilized by incubating with 37% HCl for 10 min. After washing 5 times with sterile water, the seeds were dried in a laminar flow hood, after that germinated and grown on nitrogen-free agar (Agar No.1 Oxoid) containing Hoagland solution. The seedlings were transformed by using *Agrobacterium rhizogenes* ARqua1 to generate transgenic roots.

For the nodulation experiments, *M. truncatula* cv. Jemalong with transgenic roots were transferred from agar to vermiculite (16/8 hr photoperiod at 22°C and 60% humidity) and fertilized weekly with a nitrogen-free nodulation solution for two weeks. Wild type plants were grown on nitrogen-free agar containing a nodulation solution and were inoculated directly without nitrogen-starvation. All plants were inoculated with the *S. meliloti* strain Sm2011-mRFP1 expressing a red fluorescent protein, and the nodules were harvested 2-4 weeks after inoculation.

### Transient expression in *Nicotiana benthamiana*

*N. benthamiana* was grown in a plant growth chamber at 21°C, with a 14 hr light exposure, and a10 hr dark period for 5-6 weeks. Transient transformation of tobacco leaf epidermal cells was performed with the leaf infiltration method using the A. tumefaciens strain GV3101:pMP90 at OD_600_ value of 0.05.

### Protoplast transformation

Protoplasts were isolated from *N. tabacum* BY-2 suspension cells grown at 24 °C with shaking (130 rpm) in a medium containing Murashige and Skoog salts supplemented with 30 g/L sucrose, 100 mg/L myoinositol, 255 mg/L KH_2_PO_4_, 1 mg/L thiamin-HCl, 0,2 mg/L 2,4-dichlorophenoxyacetic acid, at a pH of 5,8 and subcultured once a week. A 20 ml aliquot of a three-day-old suspension culture was centrifuged at 400 g at room temperature for 5 min and the pellet was washed with a wash-solution (0,5%(w/v) Bovine serum albumin (BSA), 0,01%(w/v), 2-mercaptoethanol, 50 mM CaCl_2_, 10 mM Na-acetate, 0,25 M mannitol, pH 5,8) and resuspended in an isolation-solution (wash-solution containing 1% cellulose R10 (Onozuka) and 0.5% macerozyme (Duchefa), pH 5.8.) and incubated at 26 °C overnight. Then, the cell suspension was centrifuged at 100 g at room temperature for 5 min, and the pellet washed once with the wash solution. The pellet was then washed again with 10 ml of w5-solution (154 mM NaCl, 125 mM CaCl_2_, 5 mM KCl, 5 mM glucose, pH 5,8-6,0) and resuspended in 5 ml of w5-solution and incubated in the dark at 4°C. The supernatant was removed, and the protoplasts were washed once with MMM-solution (15 mM MgCl_2_, 0,1% (w/v) MES, 0,5 M Mannitol, pH 5,8). The protoplasts were adjusted to 2 × 10^6^ cells/ml in MMM solution before adding ~30 μg of each plasmid DNA to 300 μl of the protoplast suspension followed by the addition of 300 μl of PEG solution (40% (w/v) PEG 4000 (Fluka), 0.4 M mannitol, 0,1 M CaCl_2_, pH 8-9) and incubated at room temperature for 10-20 min. The protoplasts were washed with w5-solution, centrifuged at 100 g for 5 min., and resuspended in 700 μl of cell culture medium containing 0,4 M sucrose. The protoplasts were then incubated in the dark for 16 hr at 26 °C.

### Immuno-localization

Semi-thin sections (60 μm) of nodule tissues were prepared using a Leica VT1000S vibratome (Leica, Wetzlar, Germany) and the sections fixed for 2 h in 4% formaldehyde in PME buffer (50 mM PIPES, 5 mM MgSO_4_, and 10 mM EGTA, pH 7.0). Then the sections were washed thrice for 10 min with PME buffer and incubated for 1 h in a blocking solution (2% (w/v) bovine serum albumin (BSA) in PBS buffer pH 7.4). Nodule sections were incubated with rabbit anti-Rab11f1 antibodies (dilution 1:20 in PBS pH 7.4 containing 0.5% (w/v) BSA) overnight at 4 °C. The samples then were rinsed three times for 10 min each with PBS and incubated with secondary goat anti-rabbit IgG Alexa Fluor 647 (Molecular Probes) (dilution of 1:50 in 0.5% (w/v) BSA in PBS) for 2 h. Nodule sections were washed thrice for 10 min each with PBS and observed under a confocal laser scanning microscope. As a negative control, thin sections of root nodules were incubated only with primary anti-Rab11F or with antirabbit antibodies.

### Fluorescence dye, BFA treatment, and microscopy

For BFA treatment, the transgenic tobacco leaves were incubated in 50 μM BFA diluted from a 50 mM stock in DMSO and then mounted on slides in the presence of BFA. The transgenic *M. truncatula* roots were inoculated with 5 μ of FM4-64 (Invitrogen, Molecular Probes) diluted from 5 mM stock in water to stain the endocytic organelles. Plant tissues and cells were observed under a confocal laser scanning microscope (Leica TCS SPE: Heidelberg, Germany) using a 63×oil-immersion objective. CLSM images were obtained using excitation/emission wavelengths at 488/500-530 nm for mGFP6, 532/ 570-620 nm for mRFP1/ Fm4-64, and 635/650-700 nm for Alexa flour 647. Images were processed using the Leica Application Suite Advanced Fluorescence (LAS AF) software.

## Results

### Localization of Rab11F in *M. truncatula* root

To study the role of Rab11F in symbiosome formation, we constructed a recombinant plasmid expressing a fusion protein of Rab11F and mGFP6. The construct was transfected into the *M. truncatula* root using the hairy root method(28), and the expression was observed using a confocal laser scanning microscope (CLSM). *M. truncatula* roots expressing GFP-Rab11F exhibited motile GFP labeled structures of uniform size, about 0.89± 0.122 (n=22) μm in diameter, randomly distributed throughout the cytoplasm (Fig 1). These structures were observed to be spherical or disk-shaped, depending on the field of view. The shape and localization patterns of these labeled structures are suggestive of the Golgi apparatus(29,30). No green fluorescence could be detected in the vacuole and the nucleus. The cytoplasm in the *M. truncatula* root tips was almost filled with GFP-Rab11F labeled punctuate structures, which moved over very small distances (Fig 1B). At the elongation zone and up to the older parts of the roots, there was a continuous decrease in the numbers per volume of GFP labeled structures, with no changes in morphology (Fig 1D). Frequently, these structures were observed moving several μm through the cortical cytoplasm before they stopped (resting phase) or even reversed their direction of movement. These observations indicated the localization of Rab11F to the Golgi apparatus.

**Figure 1:**
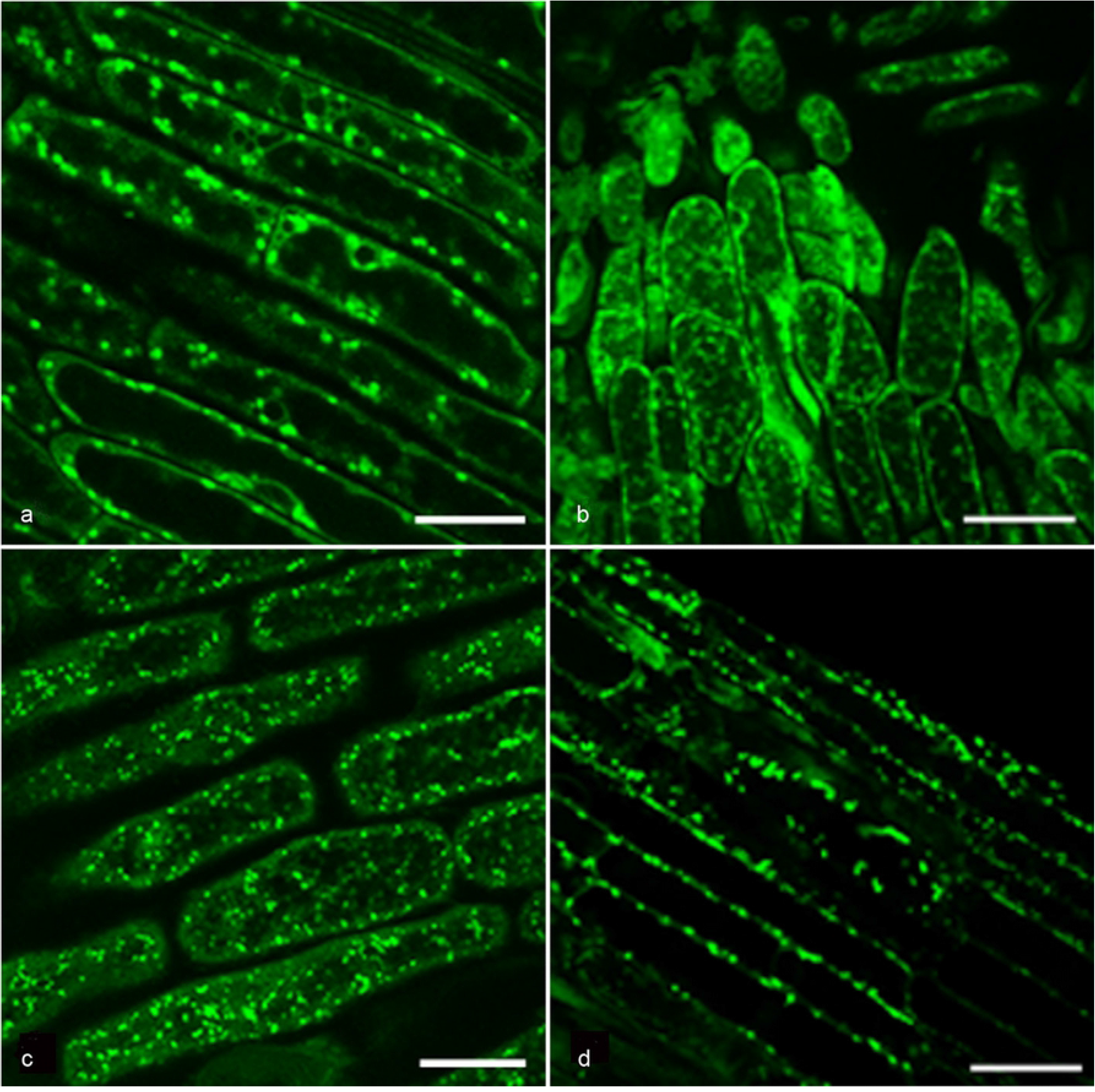
Localisation of GFP-Rab11F in *M. truncatula* root cells. Roots were transformed by the *Agrobacterium rhizogenes* strain ArQua1 carrying the GFP-Rab11F construct and investigated by confocal laser scanning microscopy (CLSM). The image shows cells in the elongation zone. In the cytoplasm of the root cells, several motile green fluorescent structures were randomly dispersed throughout the cytoplasm. Scale = 12.93 μm.

### Effect of Brefeldin A (BFA) on the morphology and streaming movement of Rab11F-mGFP6 labeled structures

To test the localization of Rab11F on the Golgi apparatus, the toxin Brefeldin A (BFA) was used. BFA is a fungal toxin which interferes with the transport of vesicles from the ER to the Golgi and alters the morphology of the plant Golgi stacks(29). Within minutes after the addition of BFA to GFP-Rab11F transiently transformed *Nicotiana benthamiana* epidermal cells, most of the GFP-Rab11F labeled structures disintegrated (Fig 2B), and the majority of the green fluorescence was dispersed throughout the cytoplasm with a higher concentration in a region around the nucleus. No fluorescence could be detected in the vacuole. The few remaining punctated structures in the cytoplasm had an average diameter of 2.2087 ±, 0.4 (n=10) larger than the GFP-Rab11F labeled structures (0.9041± 0.22 μm) (n=10) in not-treated cells. Besides, in BFA-treated cells, the streaming stop and go movement of GFP-Rab11F labeled structures was reduced to only very short distances or absent.

**Figure 2:**
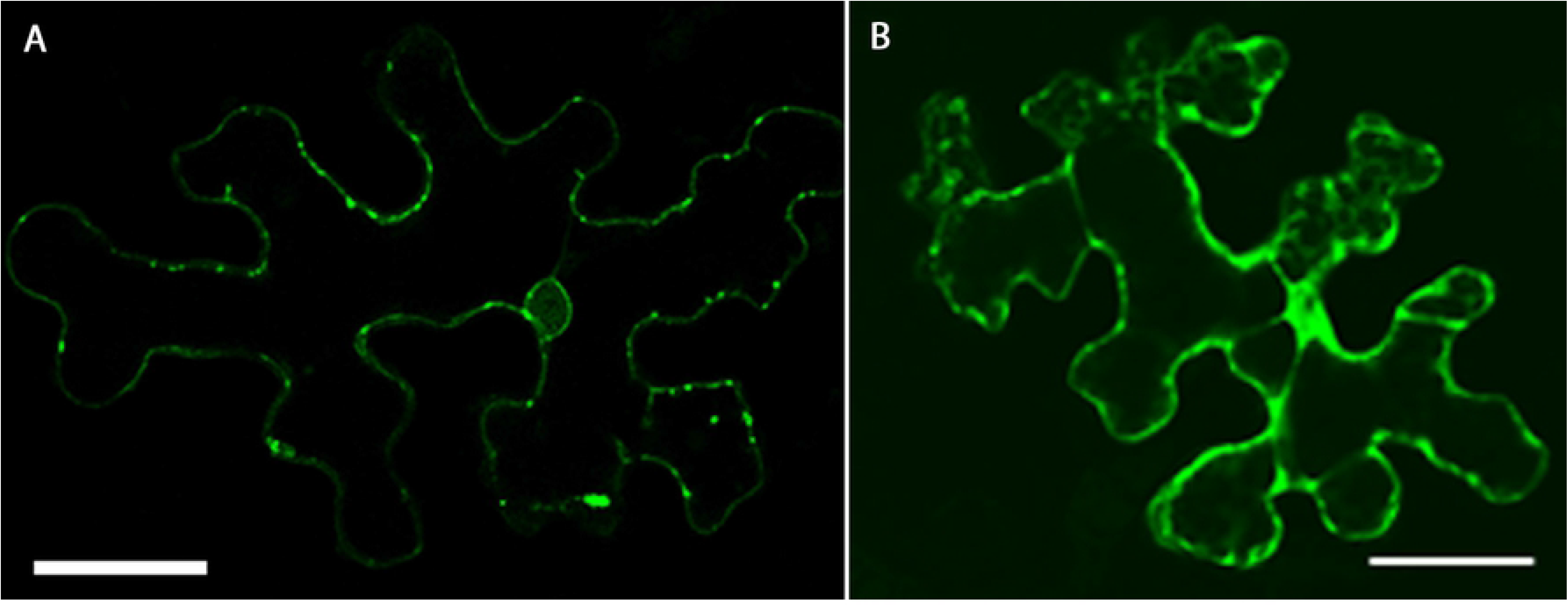
Effect of (BFA) on GFP-Rab11F expressing *N. benthamiana* leaf cells. (A) GFP-Rab11F expressing leaf cell without BFA treatment. (B) Leaf epidermal cell 30 min after application of BFA (50 μg/ml). The GFP-Rab11F labeled structures formed non-motile aggregates. Scale = 25 μm.

### Effect of dominant–negative Rab11F mutant on the Rab11F labeled structures

To study the function of Rab11F, an inactive mutant (GDP-locked) Rab11F, Rab11F(S29N), was generated by site-directed mutagenesis. Despite numerous transfection experiments, we were unable to transfect the Rab11F(S29N) construct into *M. truncatula* roots using the hairy root transformation method. However, transfection of the recombinant Rab11F(S29N) construct into tobacco BY-2 protoplasts was successful, and we observed Rab11F(S29N)-GFP evenly distributed in the cytoplasm (Fig 3C) without any specific localization as seen with the wild type GFP-Rab11F (Fig 3A). These results suggested that Rab11F is vital in *M. truncatula* cells, and its loss of function leads to cell death.

**Figure 3:**
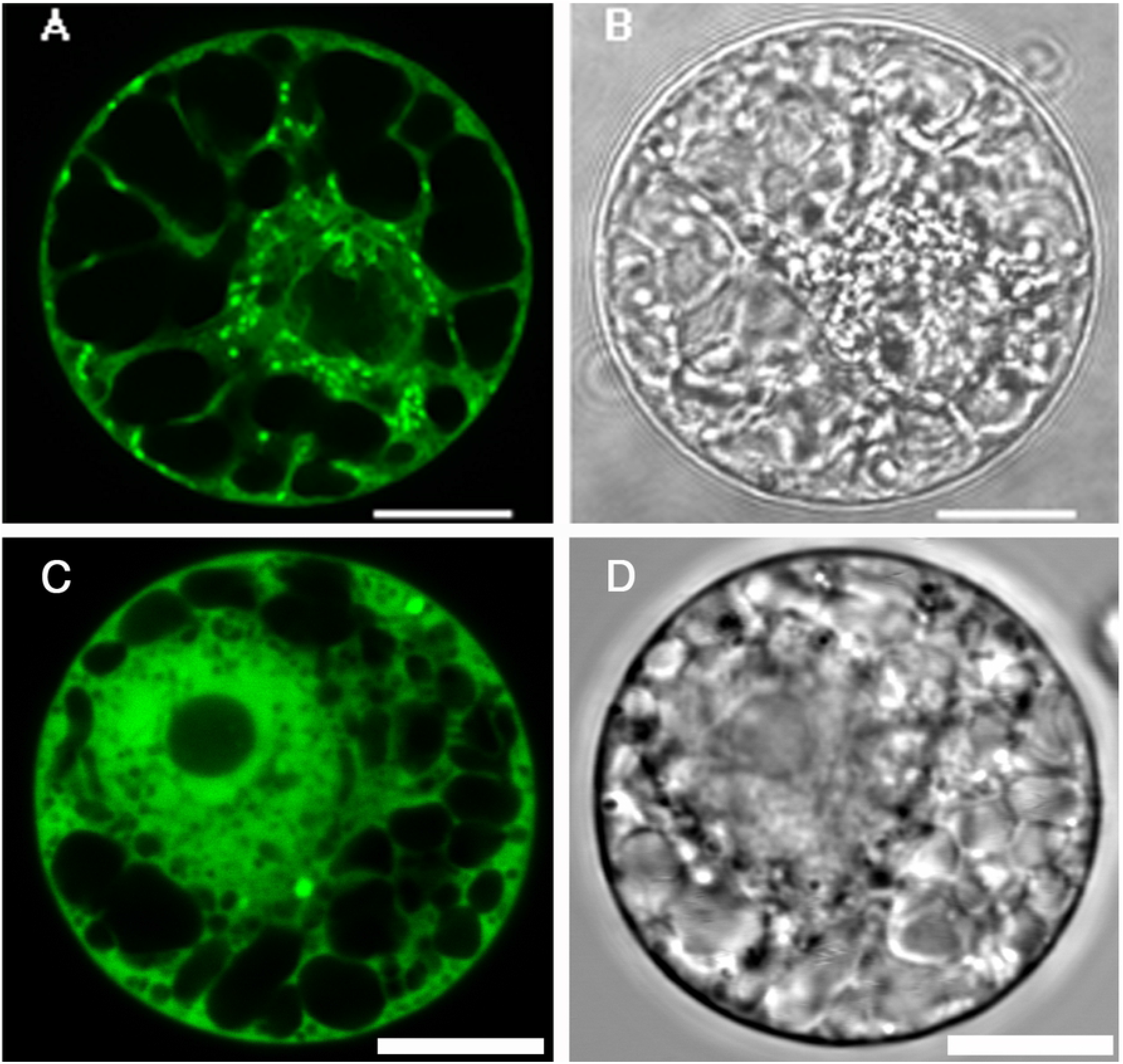
Localization of GFP-Rab11F and dominant negative mutant GFP-Rab11FS29N in tobacco BY2-protoplasts: (**A-B)** The images show tobacco By-2 protoplasts expressing GFP-Rabf11F green punctuate structures could be observed and were randomly dispersed throughout the cytoplasm. **(C-D)** The images show tobacco BY-2 protoplasts expressing the GFP labeled dominant negative mutant Rab11F-S29N. The GFP fluorescence was homogenously distributed throughout the cytoplasm. Several protoplasts transfected with the mutant showed signs of ongoing cell death. Scale =12.84 μm.

### Intracellular localization of Rab11

To gather information on the identities of Rab11F-labeled organelles, *M. truncatula* root cells expressing GFP-Rab11F were incubated with FM4-64 for a short (30 min) and a long time (2 h). FM4-64 is a lipophylic dye used for staining endocytic vesicles and the vacuolar membranes. It acts in a time-dependent manner moving with the endocytotic pathway from the plasma membrane to vesicles and finally to the vacuole membrane(31). The plasma membrane of the *M. truncatula* root cells was stained immediately after the application of FM4-64 (Fig 4B). A few GFP-Rab11F labeled structures were rapidly stained after 5 minutes. (Fig 4C arrowheads). Thirty minutes after the FM4-64 application, some small GFP-Rab11F labeled organelles (diameter of 0.320 ± 0.03, n=10) were additionally stained (Fig 4E), but these were the exceptions, and the red and green fluorescence of most structures were separated from each other. After two hours of incubation, the GFP-Rab11F labeled structures were completely stained (Fig 4I), and concomitantly the level of the FM4-64-staining of the plasma membrane was reduced.

**Figure 4:**
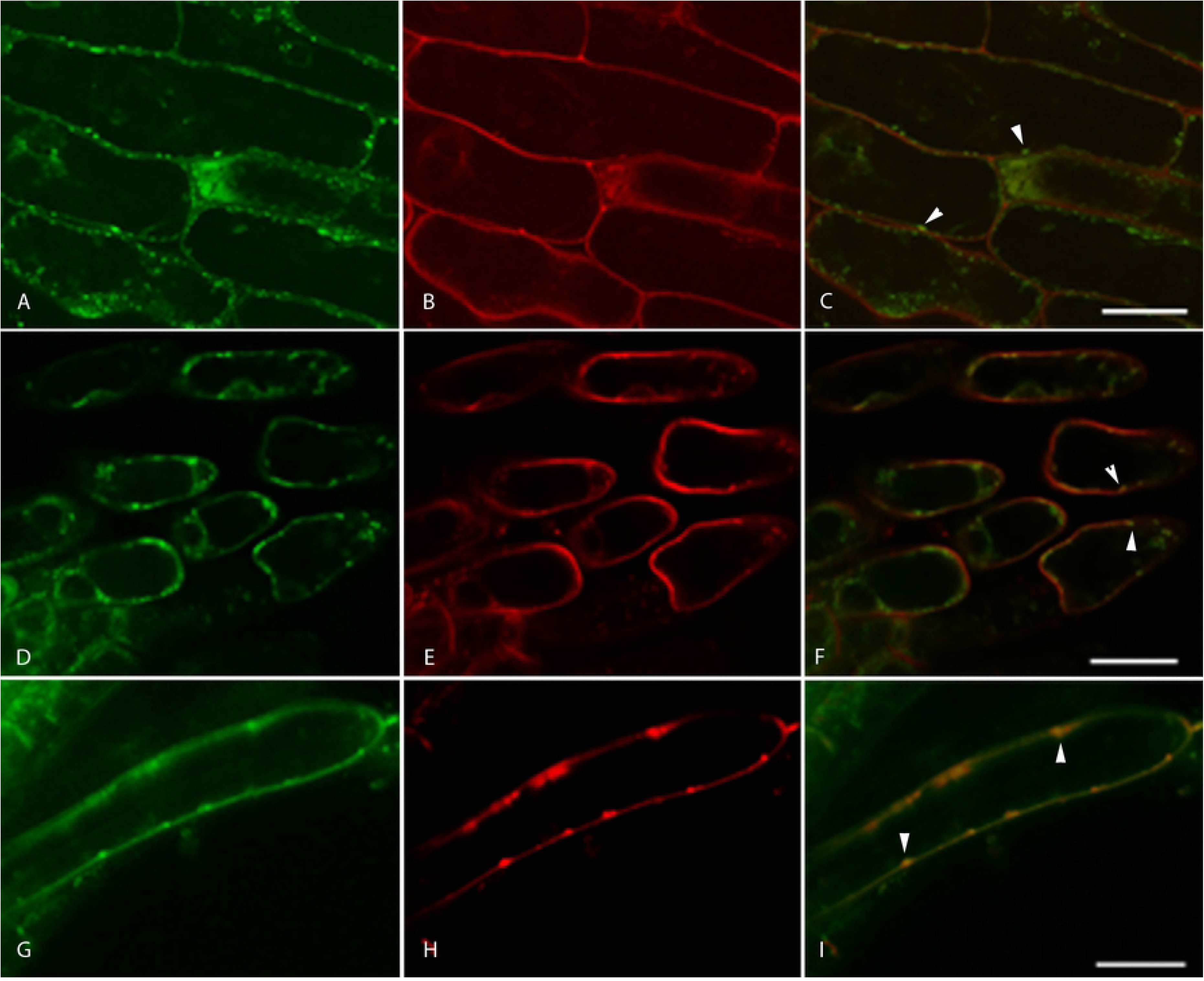
Colocalization of GFP-Rab11F and FM4-64 in *M. truncatula* root cells. Images were taken immediately after application of (A-C), after 30 min (D-F) and after 2h (G-I). GFP channel (A, D, G), dsRed channel (B, E, H) and overlay (C, F, I). Directly after application of FM4-64 the plasma membrane was stained. After 30 min GFP-Rab11F labeled structures were partly stained by FM4-64. After 2 hours, FM4-64 also stained all structures, which were labeled by GFP-Rab11F. Scale = 15 μm.

### Rab11F is located to the trans-Golgi network

For the further elucidation of the type of structure(s) labeled by GFP-Rab11F, we performed a co-localization experiment using GmMan1 fused to mCherry protein and GFP-Rab11F. GmMan1 is an α-1,2-mannosidase-I from soybean that localizes at cis-Golgi stacks(27). *N. benthamiana* leaves were co-transfected with the GFP-Rab11F and GmMan1-mCherry constructs. The expression showed several small motile structures within the cytoplasm in the GFP and the mCherry channels (Fig 5A and 5B). An overlay of both channels revealed that most of the Golgi bodies labeled by GmMan1-mCherry were also labeled by GFP-Rab11F (Fig 5C). A small number of structures showed either only red or green fluorescence. At higher magnification, a tricolored labeling pattern of the Golgi bodies became evident (Fig 5F). As GmMan1-mCherry is located at the cis-Golgi stacks whereas GFP-Rab11F labels the opposite side of the same structures, indicating that Rab11F is located on the trans-Golgi.

**Figure 5:**
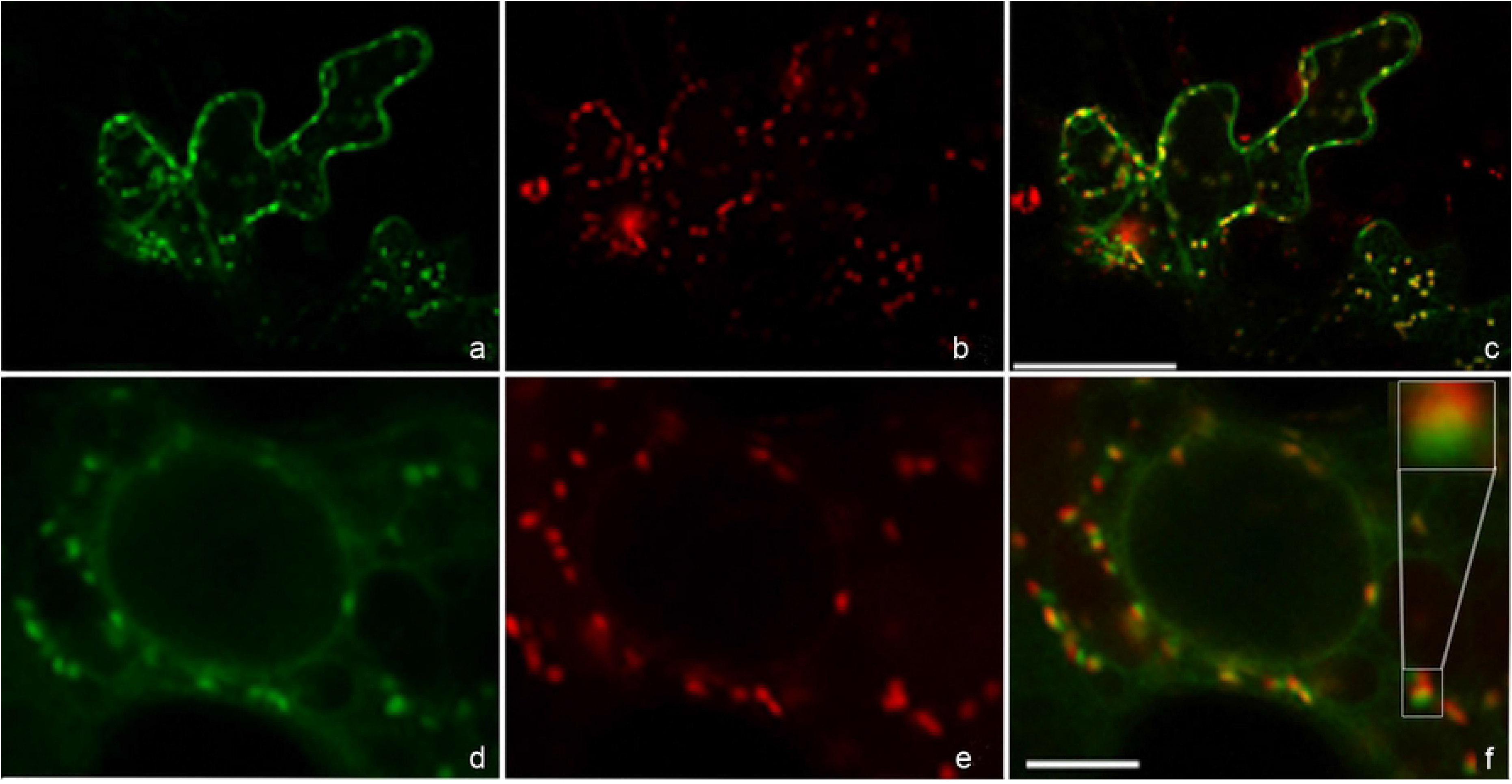
Colocalization of GFP-Rab11F and GmMan1-mCherry in *N. benthamiana* leaf epidermal cells. *N. benthamiana* leaves were co-transfected by GFP-Rab11F and GmMan1-mCherry. GFP channel (A, D), DsRed channel (B, E) overlay (C, F). GFP-Rab11F and Man1-mCherry showed a partial colocalization at Golgi bodies. (E-F) At higher magnification, the Golgi bodies revealed a tricolored labeling pattern. Scale = (A-C) 50μm, (D-F) 10μm.

### Localization of GFP-Rab11F in *M. truncatula* root nodule

Localization of GFP-Rab11F in the *M. truncatula* root nodule was determined by inoculating pBin-GFP-Rab11F transfected *M. truncatula* root cells with *S. meliloti* expressing red fluorescent protein mRFP1. Examination of nodule sections using CLSM revealed weak GFP-fluorescence on numerous punctate structures (data not shown). Most likely, the ripening of the GFP was hindered due to the low level of free oxygen in root nodules. Thus, we decided to use a peptide-specific polyclonal anti-Rab11F antibody(26) to investigate Rab11F-localization. Thin sections of 4-6 week-old, transformed root nodules inoculated with *S. meliloti* expressing mRFP1 were fixed, and immune-staining was carried out using an Alexa647-conjugated secondary antibody and examined using CLSM. The same procedure was repeated with non-transformed root nodules of wild type *M. truncatula* and *M. sativa*. Controls using the pre-immune serum or without primary antibody showed no label at any membrane.

In transgenic root nodules, anti-Rab11F-antibodies labeled small structures of nearly uniform size (1 μm) randomly dispersed in the cytoplasm especially around the infection thread, but showing high accumulation at the thread tip, where most of the Rab11F-labeled green structures were smaller (0.5 μm) (Fig 6A). Rab11F was also located on ring-like structures near the infection thread membrane (Fig 6A-6C), which were probably formed by the fusion of Rab11F-labeled structures. Interestingly, Rab11F-labeled structures also accumulated near putative release sites or infection droplets as revealed by the enlargement of the infection thread (Fig 6C). In rare cases, bacteria were not released from the infection droplet but directly from the infection thread where they pass the cell-cell border and enter the next cell (Fig 6D-6F). During the release, the bacteria were covered by Rab11F-labeled structures.

**Figure 6:**
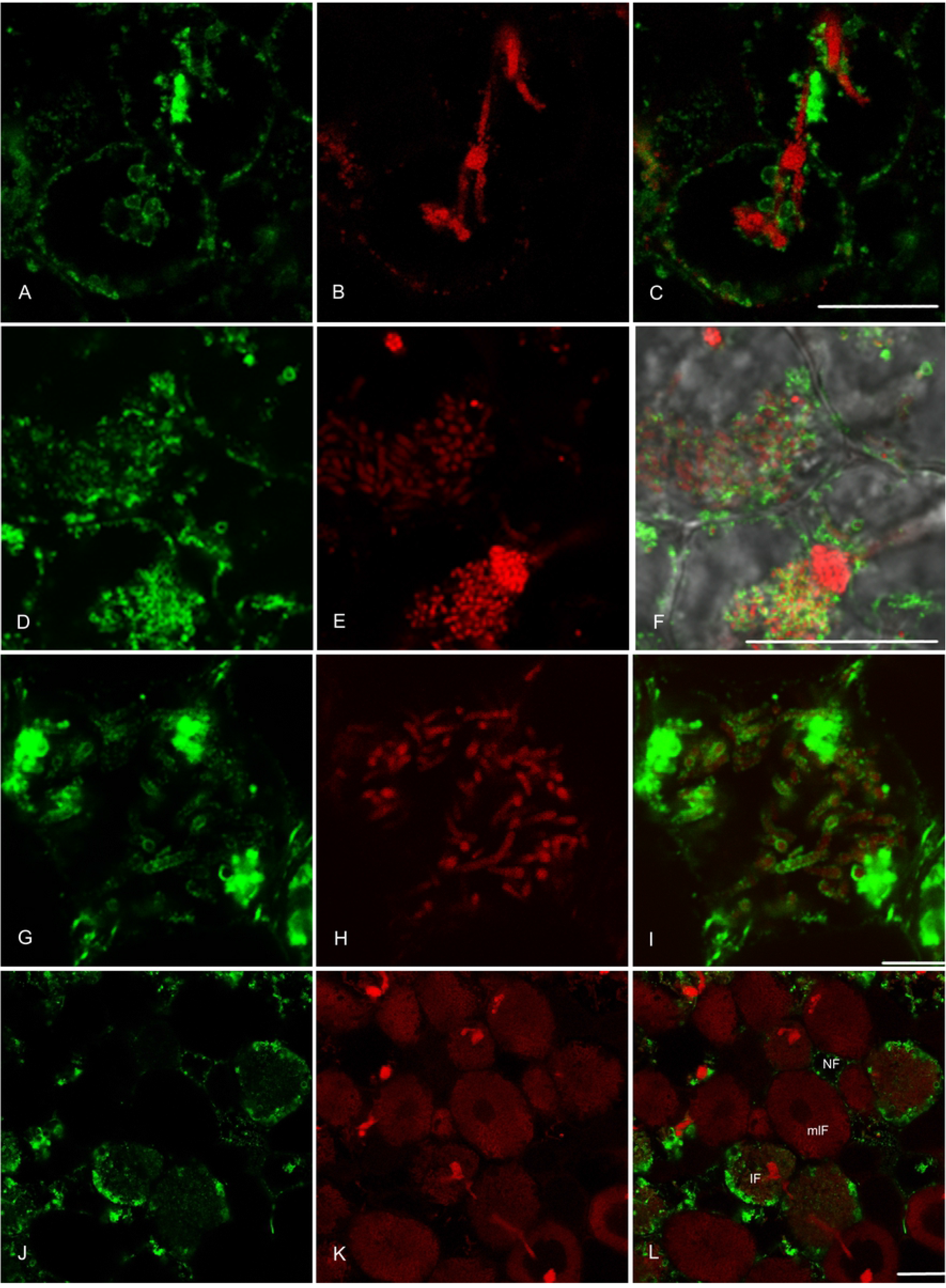
Localization of Rab11F and *S. meliloti*-mRFP1 in the young and mature root nodule cells. **(A-C)** Rab11F-labeled structures accumulated around the infection thread and at the tip. Some of them were located on the extension of the infection thread membrane, forming a ring. The infection thread was enlarged to become an infection droplet with a diameter of 10 μm and surrounded by Rab11F-positive structures. **(D-F)** At a cell-cell border, the infection thread membrane had fused with the plasma membrane of the neighbor host cell; the bacteria were then released into the cytoplasm. **(G-I)** Localization of Rab11F in young infected cells. The infected cell contained several bacteria enclosed by a membrane labeled by Rab11F. Numerous Rab11F labeled green structure were located on the symbiosome membrane. **(J-K)** Localization of Rab11F in matured infected cells (mIF). The fully differentiated infected cells (mIF) were filled with bacteroids; however; no Rab11F positive structures are detectable. Most of the bacteroids were no longer enclosed by membrane labeled by Rab11F in matured infected cells (mIF). The young infected cells (IF) contained some Rab11F positive structures surrounding the bacteroid. In non-infected cells (NIF), numerous small punctate structures were dispersed in the cytoplasm with a diameter of approximately 1 μm. Scale =(A-C) 25μm., (D-F) 25μm, (G-I) 7.5μm, (J-L)=10μm.

After the invasion, rhizobia begin to differentiate into nitrogen-fixing bacteroids. Each bacteroid was found to be enclosed by a membrane, which was labeled by anti-Rab11F antibodies (Fig 6G-I). In young infected *M. truncatula* cells, bacteroids were of different sizes: some were rod-shaped and 1 μm in diameter while others were enlarged to a length of ~ 5 μm, 5 times larger than non-differentiated rhizobia.

During the maturation of the infected cells, the amount of Rab11F-labeled structures steadily decreased. There were cells not filled with bacteroids, and in between the bacteroids were numerous small Rab11F-labeled structures enclosing some of them, but there were some bacteroids that were not surrounded by Rab11F-positive membranes (Fig 6J-6L). In mature nitrogen-fixing cells, some Rab11F-labeled structures could be detected between the bacteroids, but the majority of them were no longer surrounded by the Rab11F-labeled membrane (Fig 6J-L).

## Discussion

The engulfment of *Rhizobia* into plant host cells occurs when the infection thread reaches the target cell. Within the host cell, each invaded bacterium is bound by the host-derived symbiosome membrane. As the bacteria divide, the symbiosome membrane surface area also has to expand, in most cases almost a hundred-fold in comparison to the original plasma membrane-derived symbiosome membrane(2). Thus, a large amount of membrane material must be mobilized to allow the expansion of the symbiosome membrane.

Rab11F, a regulator of membrane trafficking, is highly expressed in nodules when the symbiosome membrane proliferates(26). We postulate that Rab11F is involved in the expansion of the symbiosome membrane. This notion is further supported by the findings that the high level of expression of the *rab11f* gene is correlated with that of the *dnf1* gene, which is essential for the establishment of symbiosis(23,26). To test this hypothesis, we traced the presence and the intracellular distribution of Rab11F by transfecting *M. truncatula* with a plasmid construct expressing a GFP-Rab11F fusion protein. Rab11F was shown to be localized on punctate structures in a pattern similar to that of the Golgi apparatus under normal conditions. By treatment with Brefeldin A (BFA), a fungal inhibitor causing aggregation of TGN and endosomes(32), the Rab11F labeled structures were aggregated from the spherical structure to BFA-induced compartments, which had been seen in BFA-treated TGN and endosomes indicating that these punctate structures are either TGN or endosomes(33,34). Co-localization experiments using the cis-Golgi marker, GmMan1, showed three colored organelles, red, yellow, and green. Rab11F labeled structures were located opposite to the cis-Golgi, indicating that Rab11F was located on TGN. However, there were some of these structures showing only one color; green, or red, without any other colors observed. These occurred possibly through the movement of the labeled structures showing only one side of them. The presence of Rab11F at the TGN is consistent with the function of Rab11-subfamily in animals and yeast, where Rab11s are involved in the recycling of plasma membrane-derived endosomes(19,20) and the secretion of TGN buddings to the plasma membrane(22). In plants, several studies have shown that Rab11/RabAs are associated with the secretory pathway from the TGN to the plasma membrane(35–37).

In animal cells, TGNs are the main sorting station of the post-Golgi pathway, delivering various cargo to the plasma membrane, and receiving cargo from endosomal compartments(38,39). In plants, some reports showed that TGN was similar to the partially coated reticulum (PCR) identified as an early endosomal compartment containing clathrin-coated pits(40). By electron microscopy/tomography of non-meristematic cells revealed two types of TGNs: GA-TGNs (Golgi associated TGN) and GI-TGNs (Golgi released independent TGNs)(41,42). GA-TGNs are located on the trans-side of the Golgi apparatus, whereas GI-TGNs are located distantly from Golgi and move independently. GI-TGNs are smaller than GA-TGNs and mostly fragment into SVs and clathrin-coated vesicles (CCVs)(43). In Maize, several secretory vesicles fragmented from GI-TGN were observed in the vicinity of the growing point of the secondary cell wall or called wall in growth (WIG) of the basal endosperm transfer cell (BETC) during the maturing process to supply of new cell wall polysaccharides from the Golgi(44).

In the *M. truncatula* nodules, various Rab11F-labeled structures were found dispersed throughout the cytoplasm at high density and presented the same pattern as GA-TGN. However, several Rab11F labeled structures were located in the vicinity of the infection thread, and the peribacteroid membrane displaying smaller in size (~ 0.5 μm) suggesting that they are GI-TGN or TGN derived vesicles. These interpretations are consistent with previous electron microscopic studies in which numerous Golgi-derived vesicles were found in the vicinity of the infection thread membranes(45). Rab11Fs were located on both the infection thread membrane and the peribacteroid membrane, even though the structure of the peribacteroid membrane are different from those of the infection thread membrane, which generates the cell wall.

When the infection thread reaches its target cell, the CLSM–images revealed two locations where the bacteria are released, namely, as infection droplets or into intercellular space. The bacterial invasion process in the intercellular space occurs through the formation of an extension of the infection thread projecting into the underlying cell layer through the fusion with the distal cell wall, thereby allowing bacteria to enter the intercellular space and to degrade the proximal cell wall. This invagination is similar to that seen at the beginning of the root hair curl(46). However, upon releasing the bacteria, the formation of the infection thread wall is suppressed, allowing the bacteria to make contact with the host cell plasma membrane, resulting in the engulfment of the bacteria. The symbiosome membrane acquires Rab11F immediately during the invasion process and throughout its differentiation. When the bacteroids matured, no Rab11F was present on their peribacteroid membranes.

In animals, bacteria enter the cell through phagocytosis and undergo a maturation process within the endosome, from the Rab5-labeled early endosome to the Rab7-labeled late endosome(9). However, in plants, symbiosomes do not acquire Rab5 at any stage of bacterial development but do so after the bacteria have stopped dividing (12). The presence of Rab11F on the TGN and the peribacteroid membrane during bacterial engulfment and symbiosome maturation suggests that the function of Rab11F is involved in secretion, which is consistent with reports of the presence of membrane-type syntaxin SYP132. SYP132 resides on organelles of the secretory system and on symbiosome membranes (12,47), suggesting that the secretory pathway is essential for the symbiosome formation. This is in agreement with the report of Wang et al. (2010) postulating that effective symbiosome formation requires an orderly secretion of protein constituents through coordinated up-regulation of a nodule-specific pathway involving GTPase Rab11(23). In bacterial infections of humans and animals, Rab11 is a prominent target for the invading pathogen (48). As shown by Limpens et al. (2009), the symbiosome membrane does not recruit the TGN marker SYP4 and is therefore not part of the conventional secretory pathway. The symbiosomes are locked in an SYP132/Rab7-positive endosome stage(12).

Further on, two highly homologous exocytotic vesicle-associated membrane proteins (VAMPs) are required for the biogenesis of the symbiotic membrane(49). Silencing of VAMP72 blocks the rhizobial symbiosome formation as well as arbuscule formation in mycorrhizal interaction. This suggests that an ancient exocytotic pathway forming the periarbuscular membrane compartment has also been coopted in the Rhizobium–Legume symbiosis(49). A symbiosome-specific modification of the extracellular matrix, namely pectins, could be part of the evolution of this specific pathway(50). In this complex network of changing membrane identities Rab11F, and specifically the analyzed isoform RabA6b, could contribute to delivering specific cargos from the TGN to the early symbiosome.

Most interestingly, *Rhizobial* release occurs only in the very young plant cells close to the meristem(51). Cell division involves the endocytotic uptake of material from the apoplastic space (52). A key regulator of this process is the Rab11 isoform RabA1d(18). This fact could give a clue to the question of the evolutionary origin of the intracellular symbiosis. Ancestral forms of *Rhizobium* could have gained access to the endocytotic pathway needed for cell plate formation in dividing cells. Plant mutants or active interference of Rhizobia could have contributed to the establishment of the symbiosome as a transient organelle.

## Acknowledgments

This work was supported by grants from the DFG and Bielefeld University. The authors thank Prof. Dr. Ton Bisseling for the S. meliloti strain Sm2011-mRFP1, Dr. Andreas Nebenführ for the GmMan1-mCherry vector, and Prof. Dr. Prapon Wilairat for a critical reading of the manuscript.

